# On the independent loci assumption in phylogenomic studies

**DOI:** 10.1101/066332

**Authors:** W. Bryan Jennings

**Affiliations:** Departamento de Vertebrados, Museu Nacional, Universidade Federal do Rio de Janeiro, Rio de Janeiro, RJ, 20940-040, Brazil.

## Abstract

Studies using multi-locus coalescent methods to infer species trees or historical demographic parameters usually require the assumption that the gene tree for each locus (or SNP) is genealogically independent from the gene trees of other sampled loci. In practice, however, researchers have used two different criteria to delimit independent loci in phylogenomic studies. The first criterion, which directly addresses the condition of genealogical independence of sampled loci, considers the long-term effects of homologous recombination and effective population size on linkage between two loci. In contrast, the second criterion, which only considers the single-generation effects of recombination in the meioses of individuals, identifies sampled loci as being independent of each other if they undergo Mendelian independent assortment. Methods that use these criteria to estimate the number of independent loci per genome as well as intra-chromosomal “distance thresholds” that can be used to delimit independent loci in phylogenomic datasets are reviewed. To compare the efficacy of each criterion, they are applied to two species (an invertebrate and vertebrate) for which relevant genetic and genomic data are available. Although the independent assortment criterion is relatively easy to apply, the results of this study show that it is overly conservative and therefore its use would unfairly restrict the sizes of phylogenomic datasets. It is therefore recommended that researchers only refer to *genealogically* independent loci when discussing the independent loci assumption in phylogenomics and avoid using terms that may conflate this assumption with independent assortment. Moreover, whenever feasible, researchers should use methods for delimiting putatively independent loci that take into account both homologous recombination and effective population size (i.e., long-term effective recombination).

## Introduction

A key assumption of phylogenomic studies that use multi-locus coalescent methods to estimate species trees and historical demographic parameters such as effective population sizes, population divergence times, and gene flow holds that each DNA sequence locus is “independent” from other sampled loci. This assumption is important because genealogical histories (i.e., gene trees) of sampled loci are considered as true replicate samples depicting the ancestry of a genome in these statistical analyses (Edwards & Beerli 2000; Arbogast et al. 2002; Wakeley 2009). Indeed, the property of genealogical independence of loci confers benefits to phylogenomic studies because larger numbers of independent loci enhance the accuracy and precision of parameter estimates (Pluzhnikov & Donnelly 1996; Edwards & Beerli 2000; Arbogast et al. 2002; Jennings & Edwards 2005; Felsenstein 2006; Lee & Edwards 2008; Smith et al. 2013; Costa et al. 2016). Although the independent loci assumption is often mentioned in coalescent-based studies, there is significant variation in how this assumption has been phrased and interpreted.

We will now examine some examples taken from the literature, which show how researchers have treated the independent loci assumption in phylogenomics (italics and bold are mine). Arbogast et al. (2002) wrote: “*Indeed, the variance associated with estimates of divergence time between recently diverged species can be minimized not by sequencing a large number of sites per locus but by sequencing a large number of **independently segregating loci***;” Hudson & Coyne (2002): “*For results concerning multiple loci, we assume **statistical independence of the gene trees at different loci***;” Yang (2002): “*It is assumed there is no recombination within a locus and **free recombination between loci***;” Hey & Nielsen (2004): “*A key assumption of the method is that the **locus being studied has been evolving neutrally and that it has been drawn at random from all loci, with respect to genealogical history***;” Bryant et al. (2012): “*The **genealogies for separate markers are conditionally independent** given the species tree*;” McCormack et al. (2012): “*Although it is increasingly feasible to sequence entire genomes, identifying portions of the genome that are orthologous and **independently sorting** is highly desirable from the perspective of analyses that take coalescent stochasticity into account*;” Reilly et al. (2012): “*Our demographic parameter estimates may depend on the assumptions of the IM model, which include loci **independently assort in meiosis***;” and lastly, O’Neill et al. (2013) stated “*To maximize coverage of the genome and independence of loci, we chose loci that ranged from approximately 200-650 bp in length, **were widely distributed across all 14 linkage groups and were on average about 50 cM from other included loci** on the* Ambystoma *linkage map*.” As this brief survey shows, researchers have identified independent loci in at least two different ways. In the first, independent loci are those that have independent genealogical histories, whereas in the second independent loci are those that undergo Mendelian independent assortment in meiosis. Several of the above bold-emphasized excerpts including “independently segregating loci,” “free recombination between loci,” “independently sorting,” and loci being “50 cM from other included loci,” presumably also refer to loci that undergo independent assortment. A pair of intra-chromosomal loci that are separated by a map distance of at least 50 centimorgans (cM) are generally considered to be independently assorting in meiosis with respect to each other. Thus, the independent loci assumption—as used in phylogenomic studies—has evidently been conceptualized in at least two different ways. Studies that refer to loci with independent genealogies are correctly encapsulating the independent loci assumption in phylogenomics, whereas other studies are apparently confusing this assumption with the independence assumption used in classical Mendelian genetics. However, it is unclear whether the alternative interpretation (i.e., “independent assortment”) can also satisfy the independence assumption in phylogenomics. Clarification of this inconsistency is important otherwise the potential exists for some researchers to use incorrect or inefficient criteria for identifying independent loci.

In order to precisely differentiate these two interpretations of the independence assumption, we can think of each as a specific criterion: the first (hereafter criterion 1), considers loci to be independent of other sampled loci if their genealogical histories are effectively independent of each other, whereas under the second (hereafter criterion 2), sampled loci are independent of each other if they undergo independent assortment. Criteria 1 and 2 are equivalent when considering two loci found on different chromosomes—just as loci found on different chromosomes will undergo independent assortment, such loci will also have independent gene trees (Wakeley 2009). However, these criteria differ from each other regarding the identification of genealogically independent loci found on the *same* chromosomes. While criterion 1 takes into account both the long-term effects of homologous recombination and effective population size (*N_e_*), criterion 2 only considers the effects of homologous recombination (i.e., no demographic component). Thus, regarding loci found on the same chromosomes, these criteria are fundamentally different from each other and this difference has important implications for phylogenomic studies.

Advances in next generation sequencing are enabling researchers to obtain phylogenomic datasets consisting of hundreds to thousands of targeted loci via in-solution sequence capture methods (e.g., Gnirke et al. 2009; Faircloth et al. 2012; Lemmon et al. 2012; Meikeljohn et al. 2016) or whole-genome sequencing (e.g., Jarvis et al. 2014). Thus, a need exists for practical methods that can identify loci that likely meet the independence assumption otherwise large genome-wide datasets may inadvertently include pseudoreplicated loci (Costa et al. 2016). One approach that has been used to identify putatively independent loci in samples has been to use complete genome data in conjunction with an *a priori* “distance threshold,” which represents the minimum intra-chromosomal “distance” between two sampled loci that are presumed to have independent gene trees. These distances have been in the form of physical distances in units of base pairs or “bp” (e.g., Sachidanandam et al. 2001; Leaché et al. 2015; Costa et al. 2016) or a recombination distance in units of cM (e.g., O’Neill et al. 2013). In other studies, researchers evaluated their datasets in an *a posteriori* manner by observing that sampled loci were separated from each other by vast intra-chromosomal distances (e.g., > 1 Mb) and therefore likely satisfied the independence assumption (e.g., McCormack et al. 2012). However, only the studies of Costa et al. (2016) and O’Neill et al. (2013) used threshold distances based on stated objective criteria: the former study implicitly invoked criterion 1, whereas the latter invoked criterion 2. Nonetheless, all studies that have made some effort to ensure that their multi-locus datasets were largely compliant with the independent loci assumption have helped move the field of phylogenomics forward.

Here, I evaluate these criteria for delimiting independent loci using empirical examples. As we will see, if sufficient data are available, then both criteria can be used to identify independent loci in a sample. However, we will also see that one of these two criteria is likely to be far too conservative for use in many phylogenomic studies.

## Materials and Methods

To illustrate the relative utility of each criterion for delimiting independent loci in eukaryotic genomes, both criteria are examined using genetic and genomic information available for the common fruit fly (*Drosophila melanogaster*) and North American Tiger Salamanders (*Ambystoma tigrinum*). Hudson & Coyne (2002) developed a theoretical framework that can be used to identify independent loci under criterion 1. These authors referred to independent loci whose gene trees are statistically independent of each other as being *independent genealogical units* or “IGUs,” which they defined as “*the number of genomic segments whose passage to monophyly is nearly independent of that for all other segments*” (Hudson & Coyne 2002). Furthermore, these authors derived a formula for estimating the total number of IGUs in a genome, which is shown here in the following general form found in Costa et al. (2016):

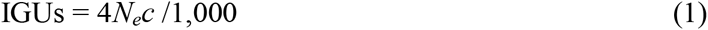

whereby *N_e_* is effective population size and the *c* is the per generation recombination rate. As mentioned earlier, criterion 1 contains a demographic component and this aspect is plainly evident in formula (1), which shows that *N_e_* plays a role in determining the number of loci with effectively independent genealogies. Thus, for a given recombination rate, large *N_e_* values translate to more IGUs per genome than smaller *N_e_* values and vice-versa. Hudson & Coyne (2002) estimated the number of IGUs in the *D. melanogaster* genome, which is based on a genetic map length of ~287 cM and *N_e_* of 10^6^ for this species (see Results and Discussion).

The number of IGUs in the *A. tigrinum* genome under criterion 1 was estimated using the genetic linkage map for the Mexican Axolotl (*A. mexicanum*), which is 5,251 cM in length (Smith et al., 2005). However, in order to use formula (1), an estimate of *N_e_* must also be supplied, which is problematic because North American Tiger Salamanders have widely varying *N_e_* depending on the species. For example, Wang et al. (2011) found that California Tiger Salamanders (A. *californiense*) had exceedingly low *N_e_* of 11-64, which may be explained by population bottlenecks or pond sizes. In contrast, Church et al. (2003), who used mitochondrial DNA, estimated the effective number of females (*N_f_*) in Eastern Tiger Salamanders (A. *tigrinum*) to be 134,000-144,000. Because autosomal loci have 4-fold higher *N_e_* than mitochondrial loci (Wilson et al. 1985), *N_e_* for autosomal loci in these salamanders are likely higher. Owing to this wide-ranging variation in *N_e_* across North American *Amybstoma* species and populations it is difficult to know which *N_e_* value should be inserted into formula (1) above. However, as these salamanders currently have a continental-wide distribution, they may have had more genetic connectivity among populations in the past. Therefore, *N_e_* values of 10^3^-10^5^ appear reasonable for our present purpose, particularly in light of the recent phylogenomic study of this entire species complex by O’Neill et al. (2013).

Criterion 2 (independent assortment) only requires a genetic linkage map for the study species or group and thus it is simpler to use than criterion 1. Thus, given the map length of ~287 cM for the *D. melanogaster* genome (Hudson & Coyne 2002), it was straightforward to estimate the number of independent loci under under criterion 2. O’Neill et al. (2013) were evidently the first researchers to use the independent assortment criterion to select their phylogenomic loci. Using the *A. mexicanum* linkage map these authors developed 95 PCR-based loci taken from all 14 linkage groups and ensured that no two loci were closer than 50 cM apart on the same chromosomes (O’Neill et al. 2013). In the current study, the total number of independent loci in the *Ambystoma* genome under criterion 2 was estimated.

## Results and Discussion

Under criterion 1, the fruit fly genome contains approximately 11,500 IGUs (Hudson & Coyne 2002). Thus, given a genome size of ~143 Mb for this species (NCBI 2016), we would expect, on average, to encounter one IGU or independent locus every ~12,500 bp along its chromosomes. However, this type of distance threshold should be regarded as a rough estimate because local recombination rates and *N_e_* vary across genomes (Costa et al. 2016). Nonetheless, this threshold value still provides us with some means for deciding whether any given nearest-neighbor pair of loci found on the same chromosome may be genealogically independent of each other or not. What are the comparable estimates under criterion 2? If we assume that loci separated by 50 cM on the same chromosomes are independent from each other, then we would conclude that there are only six independent loci in this genome. In reality, however, there must be at least seven IGUs because there must be one IGU for each of the seven chromosomes in the *D. melanogaster* genome. This means we would expect to see one independent locus per 20 Mb in the genome. In summary, the number of independent loci under criteria 1 and 2 are ~11,500 and seven, respectively, while the inter-locus distance thresholds are ~12.5 kb and 20 Mb, respectively. Clearly, criterion 2 is far too conservative to be of practical use for fruit flies.

Using equation (1), the number of IGUs in the tiger salamander genome is equal to [(4)(1,000)(5,251 cM)(0.01 cross-overs per generation)]/1,000 = 210 IGUs. If the long-term *N_e_* is instead assumed to be larger at 10^5^, then the number of IGUs increases a hundred-fold to 21,000. With these IGU estimates and knowing that the genome of *A. mexicanum* is 354 Mb in size (NCBI 2016), we can expect to see one IGU every 17 kb to 1.7 Mb depending on whether the assumed *N_e_* value is 10^5^ or 10^3^, respectively. Under criterion 2, there are 105 IGUs in the *Ambystoma* genome, which translates to about one independent locus per 3.4 Mb, on average. Although the estimated number of independent loci in the tiger salamander genome under criterion 2 is by no means a small number of loci for a phylogenomic dataset, it is still substantially smaller than the number of loci that would be obtained using criterion 1 even if a low *N_e_* were to be assumed.

The fruit fly and tiger salamander examples demonstrate that the independent assortment-based criterion for identifying genealogically independent loci is overly stringent and would therefore unfairly restrict researchers to using fewer independent loci than would be permitted under the genealogical-based criterion. Accordingly, for evolutionary studies involving multilocus coalescent analyses it is recommended that researchers use, whenever possible, the criterion of genealogical independence for independent loci (or SNPs). Although criterion 1 is more difficult to implement than criterion 2 owing to its requirement of an estimate of *N_e_*, it offers a promising approach for elucidating appropriate physical distance thresholds between independent loci in genomes. This, in turn, should allow researchers to generate phylogenomic datasets with the maximum number of genealogically independent loci or SNPs.

